# Strigolactones are chemoattractants for host tropism in Orobanchaceae parasitic plants

**DOI:** 10.1101/2022.02.17.480806

**Authors:** Satoshi Ogawa, Songkui Cui, Alexandra R.F. White, David C. Nelson, Satoko Yoshida, Ken Shirasu

**Affiliations:** RIKEN Center for Sustainable Resource Science, Yokohama, 230-0045, Japan; Division of Biological Science, Graduate School of Science and Technology, Nara Institute of Science and Technology, Ikoma, Nara, 630-0192, Japan; Department of Botany and Plant Sciences, University of California, Riverside, CA, 92521, USA; PRESTO, Japan Science and Technology Agency, Kawaguchi, Saitama, 332-0012, Japan; Graduate School of Science, The University of Tokyo, Tokyo, 113-0033, Japan

## Abstract

Parasitic plants are worldwide threats that damage major agricultural crops. To initiate infection, parasitic plants have developed the ability to locate hosts and grow towards them. This ability, called host tropism, is critical for parasite survival, but its underlying mechanism remains mostly unresolved. To characterize host tropism, we used the model facultative root parasite *Phtheirospermum japonicum*, a member of the Orobanchaceae. Here, we show that strigolactones (SLs) function as host-derived chemoattractants. Chemotropism to SLs is also found in *Striga hermonthica*, a parasitic member of the Orobanchaceae, but not in non-parasites. Intriguingly, chemotropism to SLs in *P. japonicum* is attenuated in ammonium ion-rich conditions, where SLs are perceived, but the resulting asymmetrical accumulation of the auxin transporter PIN2 is diminished. *P. japonicum* encodes putative receptors that sense exogenous SLs, whereas expression of a dominant-negative form reduces its chemotropic ability. We propose a new function for SLs as navigators for parasite roots.

## Introduction

Plant parasitism has independently evolved at least 12 times, accounting for about 1% of angiosperms or about 4500 species^1, 2^. A key trait common to parasitic plants is the ability to connect to host plants and deprive them of nutrients and water, which is often harmful to the hosts^3, 4^. Among parasitic plants, some Orobanchaceae species such as *Striga* spp. and *Orobanche* spp. are notoriously devastating invaders of major agricultural crops, leading to a multibillion-dollar economic loss annually^5, 6^. Most obligate Orobanchaceae parasites have evolved three steps to complete invasion^2^: germination near the host root in response to host-derived stimulants such as strigolactones (SLs) ^7^, active extension of the parasite’s root to host roots^8, 9^, and connection of the vasculature via formation of an invasive organ called a haustorium^2, 6, 10–12^. Although many studies have focused on germination and haustorium development, the molecular basis for host tropism is largely unknown.

*Phtheirospermum japonicum* is a facultative hemiparasite in the Orobanchaceae that completes its lifecycle with or without a host^13, 14^. With the advent of genetic tools, including ethyl methyl sulfonate-driven mutagenesis and *Agrobacterium rhizogenes*-mediated root transformation^10, 15, 16^, *P. japonicum* has been used as a model Orobanchaceae species to investigate parasitism. These studies have revealed the molecular mechanisms regulating haustorium development, vasculature connection and exchange of molecules^12–14, 16–20^. As a facultative parasite, *P. japonicum* does not require host-derived stimulants, such as SLs, for germination. Hence, we considered *P. japonicum* to be an ideal plant for studying tropic mechanisms separate from germination.

In angiosperms, α/β hydrolases DWARF14 (D14) act as SL receptors^21, 22^. After SL recognition, D14 interacts with an F-box protein MORE AXILLARY GROWTH2 (MAX2), a component of a SKP1-CULLIN-F-BOX (SCF) E3 ubiquitin ligase complex (SCF^MAX2^), to mediate SL signalling^23^. Interestingly, SCF^MAX2^ also mediates signalling driven by karrikins (KARs), a family of compounds found in smoke that regulate plant growth^24, 25^. In *Arabidopsis*, responses to KARs require *KARRIKIN INSENSITIVE 2* (*KAI2*) / *HYPOSENSITIVE TO LIGHT* (*HTL*) (hereafter *KAI2*), a paralogue of D14^26^. Whereas *KAI2* is a single gene in *Arabidopsis*, Lamiid species such as Solanales and Lamiales have duplicated *KAI2* genes. KAI2 proteins in Lamiids are categorised into three types: a “conserved” type found in most Lamiids as well as other angiosperms (KAI2c), an “intermediate” type only found in Lamiales, including parasites and non-parasites with a few exceptions such as *Orobanche* spp. (KAI2i), and a “divergent” type conserved uniquely in Orobanchaceae parasites (KAI2d)^27, 28^. Phylogenetic analyses revealed that Orobanchaceae parasites have rapidly duplicated the *KAI2d* genes, resulting in multiple genomic copies^27–29^. Some facultative Orobanchaceae parasites that do not require host-derived SLs to germinate also encode multiple KAI2d proteins, suggesting that KAI2d may have a yet uncovered function distinct from germination stimulants.

Here, we show that *P. japonicum* roots exhibited chemotropism to various SLs. Consistent with this result, *P. japonicum* roots tended to grow towards a wild-type host rather than an SL-deficient mutant host. Chemotropism to SLs was also observed in *Striga hermonthica,* but not in non-parasitic plants, suggesting that this strategy is conceivably Orobanchaceae parasite-specific. Chemotropism to SLs in *P. japonicum* is negatively regulated by ammonium in the medium, implying that parasites infect hosts when the nitrogen source is limited. We also describe a functional link between SLs and the plant hormone, auxin, in chemotropism. Thus, our findings provide a novel function for SLs as host-produced chemoattractants for parasites.

## Results

### Orobanchaceae parasitic plants exhibit chemotropism to SLs

We previously identified multiple homologues of SL receptor-encoding genes in the genomes of *P. japonicum* and other Orobanchaceae parasites^27^, although *P. japonicum* does not require exogenous SLs for germination. Therefore, we hypothesised that *P. japonicum* might use SLs as chemoattractants rather than for germination. To evaluate this putative function, we devised an experimental system in which a chemical-soaked filter paper disk was placed next to seedlings. Dimethyl sulfoxide (DMSO) at a concentration of 0.1% (v/v) was used as the negative control. Using this assay system, we demonstrated that *P. japonicum* exhibited chemotropism to three SLs (*rac*-strigol (STR), *rac*-orobanchol (ORO) and (+)-5-deoxystrigol (5DS)) and two synthetic analogues (*rac*-GR24 and Yoshimulactone Green (YLG)^30^) (Fig. 1a-f, Supplementary Fig. 1, and Supplementary Video 1). Among the tested chemicals, *rac*-STR and (+)-5DS had the strongest chemotropism-inducing activity at a concentration of 100 nM (Fig. 1g). We also compared the chemotropism activities between stereoisomers of 5DS (Supplementary Fig. 1), as the chemical structure of SLs is important for bioactivities such as germination-inducing activity and hyphal branching-inducing activity^31–33^. In contrast to (+)-5DS, the unnatural isomer (-)-5DS did not induce chemotropism (Fig. 1h). The chemoattractant activity of (+)-5DS was comparable to that of (+)-2’-*epi*-5DS, an SL found in rice^34^, and stronger than another unnatural isomer (-)-2’-*epi*-5DS (Fig. 1h). This result suggests that *P. japonicum* can efficiently sense host-derived natural SLs for chemotropic activity. In addition, using YLG, a fluorogenic agonist for SL receptors^30^, we observed asymmetrical fluorescence in the root elongation zone (Fig. 1i), indicating that asymmetrical SL recognition activity induces an asymmetrical root elongation pattern, leading to chemotropism.

**Fig. 1:**
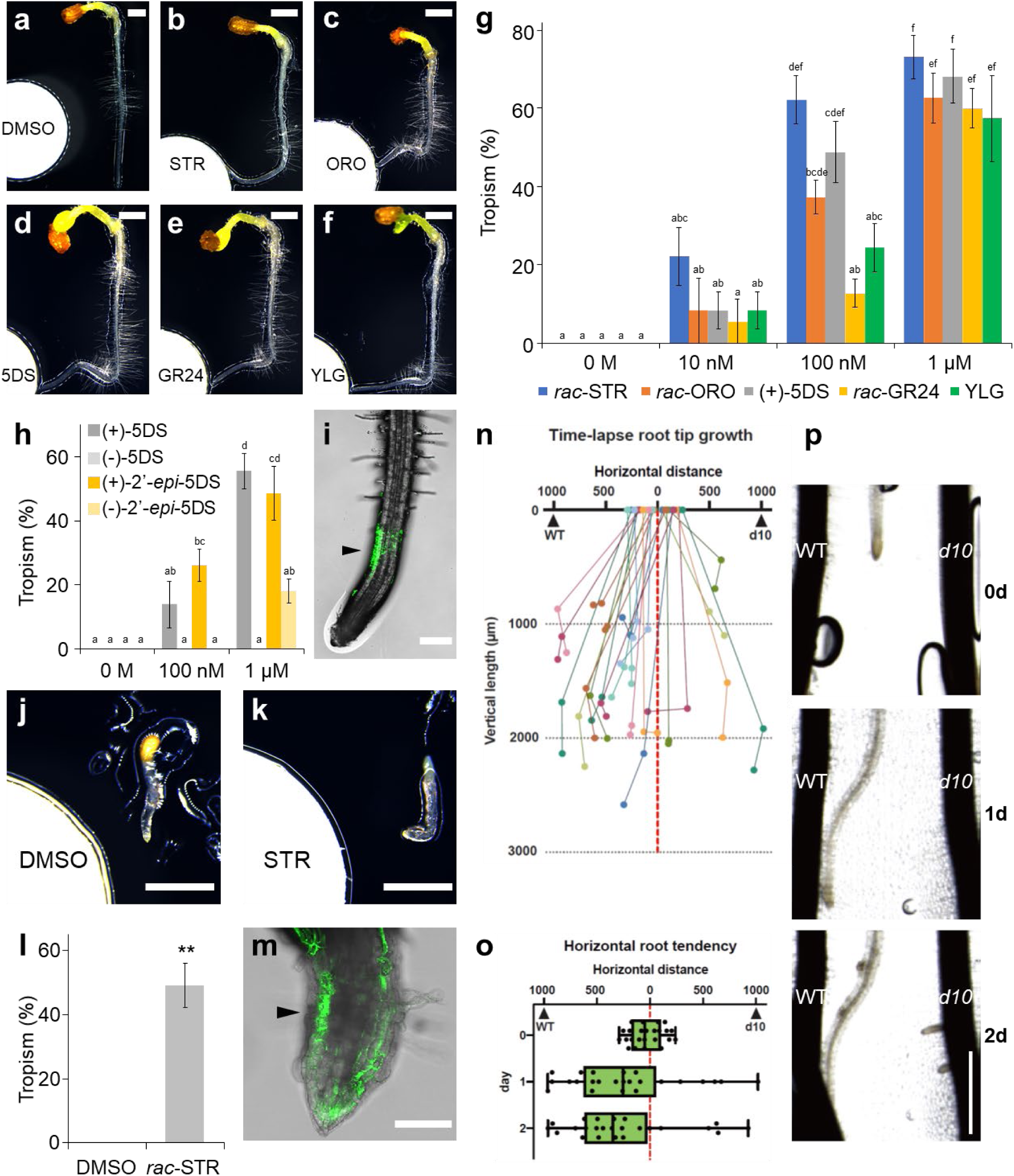
Chemotropic phenotype of Orobanchaceae parasitic plants to SLs and analogues. **a-f,** Representative images of *P. japonicum* seedlings treated with 1 µM chemical solutions diluted in 0.1% DMSO. Photos were taken 1 day after treatment. (**a**) 0.1% (v/v) DMSO; (**b**) *rac*-STR; (**c**) *rac*-ORO; (**d**) (+)-5DS; (**e**) *rac*-GR24; (**f**) YLG. **g-h** Percentage of *P. japonicum* plants that showed chemotropism to each chemical. Three independent batches (3-12 and 4-8 seedlings in each batch for **g** and **h**, respectively) for each compound. **I**, A representative image of *P. japonicum* exhibiting YLG-derived fluorescence after treatment with a 100 µM YLG solution. Filter paper disks were placed 5-mm to the left of the roots. Confocal photos were taken 6 hours after treatment. **j-k,** Representative images of *S. hermonthica* seedlings treated with 1 µM *rac*-STR or 0.1% (v/v) DMSO solutions. Photos were taken 1 day after treatment. (**j**) 0.1% DMSO; (**k**) *rac-*STR. (**l**) Percentage of *S. hermonthica* plants that showed chemotropism to 1 µM *rac*-STR. Three or four independent batches (2-13 plants) were treated with each compound. **m,** A representative image of *S. hermonthica* exhibiting YLG-derived fluorescence when treated with a 100 µM YLG solution. Filter paper disks were placed 3-mm to the left of the roots. Confocal photos were taken 24 hours after treatment. **n-p,** Root chemotropism experiments in *P. japonicum* towards intact rice. Each *P. japonicum* root was placed between a WT root and a *d10* root that were aligned vertically at approximately 2-mm apart. Quantitative measurements (n, o) and images (p) were collected at 0, 1 and 2 day(s) post co-incubation. (**n**) Time-lapse growth of. *P. japonicum* roots. Plots indicate the positions of each root. (**o**) Horizontal distances between *P. japonicum* root tips and rice roots. (**p**) Representative images of a *P. japonicum* root growing towards WT and not towards *d10*. Experiments were repeated three times with a total of twenty-four *P. japonicum* seedlings. **g,h,l** Mean ± standard error of the mean (SEM). Roots in which growth had stopped were excluded from the calculation. **i, m** Arrowheads indicate asymmetrical fluorescence. **g,h** Different letters indicate statistical significance at *P* < 0.05 (two-way ANOVA, Tukey’s multiple comparison test). **l** \*\**P* < 0.01 (Welch’s *t* test). Scale bars indicate 1 mm for **a-f, j, k, p**, 200 μm for **i**, and 100 μm for **m**, respectively

To investigate the chemoattractant activity of SLs on another parasitic plant in the Orobanchaceae family, we used *S. hermonthica* seedlings just after germination. We found that *S. hermonthica* exhibited chemotropism to *rac*-STR and had an asymmetrical YLG recognition pattern like *P. japonicum*; however, growth was limited probably due to the lack of sufficient nutrients in the typically small seeds (Fig. 1j-m). By contrast, a non-parasitic Orobanchaceae *Lindenbergia philippensis* exhibited neither chemotropism to *rac*-STR nor asymmetrical YLG recognition (Supplementary Fig. 2a,b). *Arabidopsis thaliana*, a non-parasitic plant in the Brassicaceae, also did not show chemotropism to *rac*-STR (Supplementary Fig. 2c,d). These data imply that chemotropism to SLs might be limited to Orobanchaceae parasitic plants.

Next, we conducted infection assays using *P. japonicum* and rice to test the effect of host-derived SL on host tropism. In agreement with the chemotropism results, more *P. japonicum* roots grew towards wild-type rice (WT) roots than *d10*, an SL-deficient mutant that lacks one of the SL- biosynthetic genes^35^ (Fig. 1n-p and Supplementary Video 2). Therefore, we concluded that host-derived SLs are essential for host tropism in *P. japonicum*.

### Ammonium ions are important for regulating chemotropism

Improvement of soil fertility blocks the proliferation of *Striga* spp. ^36^. Thus, we considered that tropic responses to SLs, one strategy for parasitism, may also be compromised by the presence of abundant nutrients. To test this hypothesis, we evaluated chemotropism to *rac*-STR on half-strength Murashige-Skoog (1/2MS) agar with sucrose^37^, a representative nutrient-rich medium on which *P. japonicum* can grow without parasitism^14^. In contrast to assays on water agar, chemotropism to *rac*-STR was compromised on 1/2MS agar with sucrose (Fig. 2a,b). We also showed that MS macronutrients (KH_2_PO_4_, KNO_3_, NH_4_NO_3_, CaCl_2_ and MgSO_4_) with sucrose, but not MS micronutrients, inhibited chemotropism to *rac*-STR (Fig, 2c,d and Supplementary Fig. 3). In contrast, asymmetrical YLG-derived fluorescence remained in the root elongation zone (Fig. 2e), indicating that nutrients affect the signalling pathway downstream of the SL receptors but not the perception process. To identify the component(s) that impairs chemotropism to SLs, we conducted chemotropism to *rac*-STR assays using water agar supplemented with individual MS macronutrients. Note that we added KH_2_PO_4_ to nitrogen (KNO_3_ and NH_4_NO_3_)-containing media in which the nutrient concentration was equal to 1/2MS because the addition of only a nitrogen source to these media was toxic to *P. japonicum*. We found that nitrogen, especially ammonium ions, significantly compromised the chemotropic response to *rac*-STR (Fig. 2f,g). When ammonium ions were omitted from 1/2MS, chemotropism activity was recovered, demonstrating that ammonium ions are necessary and sufficient to impair chemotropism to SLs (Fig. 2h). Next, to investigate downstream signalling, we focused on *SUPPRESSOR of MAX2 1* (*SMAX1*) genes in *P. japonicum* that encode homologues of a negative regulator of SL signalling^38, 39^. As expected, expression of *PjSMAX1* was enhanced on 1/2MS and media containing ammonium ions, suggesting that SL signalling to trigger chemotropism was suppressed (Fig. 2i-k). Overall, our data suggest that *P. japonicum* negatively regulates the SL signalling pathway in response to ammonium ions, leading to a reduction in host tropism capacity.

**Fig. 2:**
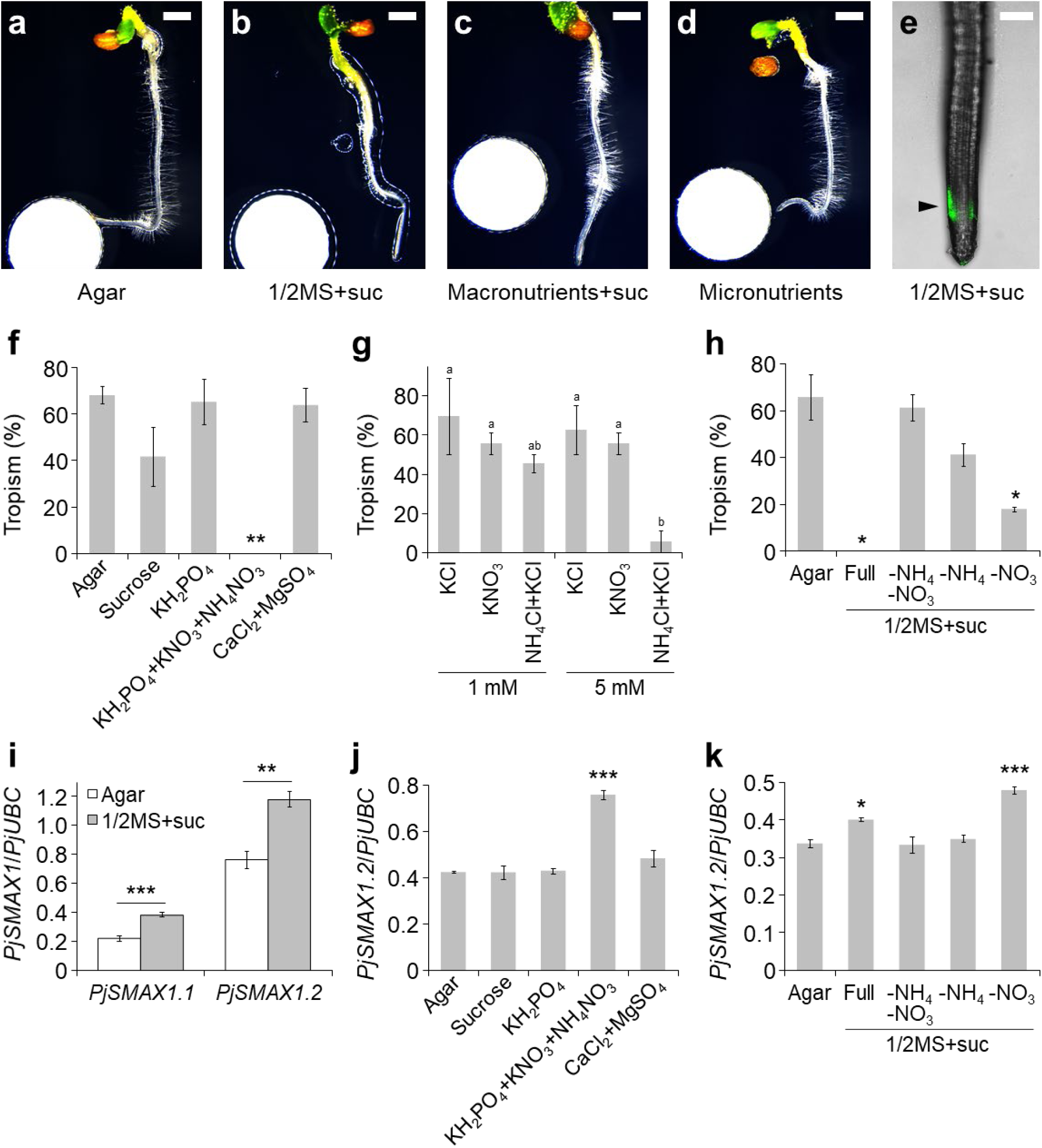
Chemotropic phenotype towards SLs on nutrient-containing media. **a-d,** Chemotropic responses to *rac*-STR in varying nutrient conditions in the presence of a 1 µM *rac*-STR solution. Photos were taken 1 day after treatment. (**a**) agar without any supplementary nutrients; (**b**) 1/2MS (Nihon Pharmaceutical) with 1% (w/v) sucrose (suc); (**c**) macronutrients of 1/2MS + suc; (**d**) micronutrients of 1/2MS without suc. **e,** A representative image of *P. japonicum* plants showing YLG- derived fluorescence when grown on 1/2MS + suc upon treatment with 100 µM YLG solution. Filter paper disks were placed 5-mm to the left of the roots. Confocal photos were taken 6 hours after treatment. The arrowhead indicates asymmetrical fluorescence. **f-h,** Percentage of *P. japonicum* exhibiting chemotropism to 1 µM *rac*-STR on each medium. Three independent batches (3 to 12 plants) were tested for each compound. Plants in which root growth had stopped were excluded from the calculation. (**f**) Assays on agar containing varying macronutrients or sucrose. (**g**) Assays on agar and a reduced nitrogen source. KCl was added to adjust the potassium concentration. (**h**) Assays on 1/2MS + suc agar with a limited nitrogen source. **i-k**, Relative expression level of *PjSMAX1* determined by RT-qPCR. Representative data are shown using *PjUBC2* as the reference gene. Experiments were performed three times with similar results. (**i**) Expression levels of *PjSMAX1.1* and *PjSMAX1.2* on agar without nutrients or 1/2MS + suc (4 technical replicates). (**j**) Expression level of *PjSMAX1.2* on agar with each macronutrient or sucrose (4 technical replicates). (**k**) Expression level of *PjSMAX1.2* on 1/2MS + suc agar with a limited nitrogen source (3 technical replicates). **f-k** Mean ± SEM. **f, h, j, k** \**P* < 0.05, \*\**P* < 0.01, \*\*\**P* < 0.001 (Welch’s *t* test) in comparison with the no nutrient treatment. **g** Different letters indicate a statistical significance at *P* < 0.05 (two-way ANOVA, Tukey’s multiple comparison test). **i** \*\**P* < 0.01, \*\*\**P* < 0.001 (Welch’s *t* test). Scale bars indicate 1 mm for **a-d** and 200 µm for **e**, respectively.

### Auxin response contributes to the tropism to host

Auxin is a phytohormone well known to regulate tropisms, as exemplified by gravitropism^40^. To test whether an auxin response also regulates chemotropism to SLs, we used *P. japonicum* hairy roots transformed with the 3X mCherry-nuclear localisation signal (NLS) module driven by the auxin- responsive promoter *DR5*^12, 19^. Asymmetrical activation of the auxin response in root tip epidermal cells was observed in the presence of *rac*-STR but was diminished on 1/2MS medium (Fig. 3a-h). This STR- and starvation-inducible asymmetrical auxin response coincided with the chemotropism-exhibiting condition. We further focused on PIN-FORMED2 (PIN2), one of the PIN-family auxin efflux transporters^41^, as PIN2 plays a major role in root tropisms ^42^ and PjPIN2 is a representative PIN localised in *P. japonicum* root epidermal cells^19^. Using the PjPIN2-Venus module driven by the native promoter, we found an asymmetrical increase in PjPIN2 accumulation in epidermal cells of the root elongation zone in water agar (Fig. 3i, j, m, n), but not in a medium containing 1/2MS+suc (Fig. 3k, l, o, p). These results suggest that local SL perception leads to local auxin accumulation for tropism, likely via PIN2, and that the nutrient-based inhibition of chemotropism occurs between SL perception and PIN2 accumulation.

**Fig. 3:**
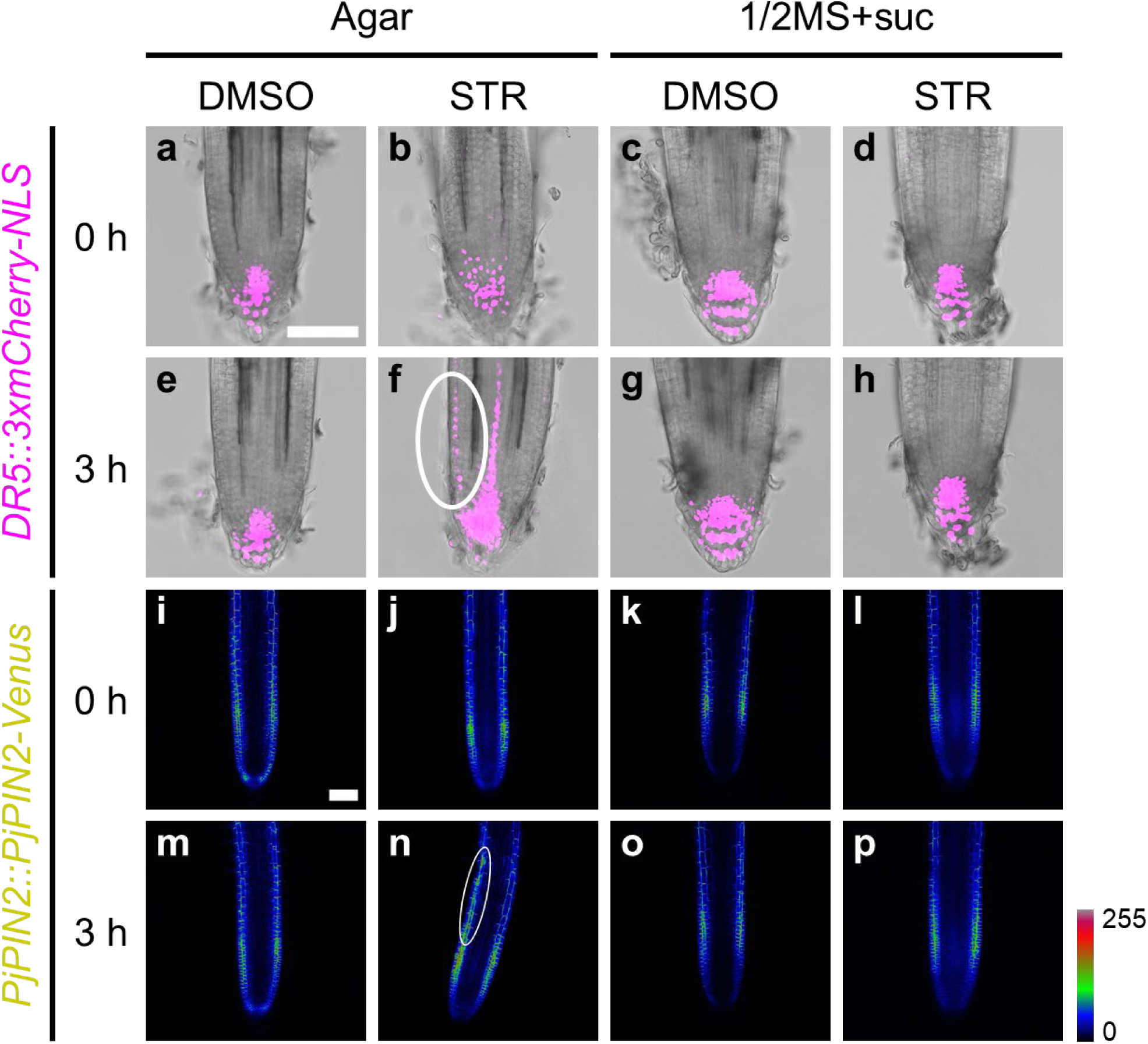
Expression dynamics of auxin-related genes upon SL treatment. Expression patterns of the *DR5* (**a-h**) and *PjPIN2* (**i-p**) promoters driving expression of a fluorescent marker gene upon treatment with 1 µM *rac*-STR or 0.1% (v/v) DMSO solutions. Filter paper disks were placed 5-mm to the left of the roots. Confocal photos were taken at the indicated time points after treatment on media with (1/2MS+suc) or without (agar) nutrient. Bright-field and fluorescent images were merged in **a-h**. Venus fluorescence intensity was represented in a Rainbow RGB spectrum in **i-p** using ImageJ^66^. White ovals indicate asymmetrical expression. The scale bars in **a** and **i** also refer to **b-h** and **j-p**, respectively. Bars = 100 µm.

### KAI2d proteins are the SL receptors involved in chemotropism

Next, we investigated if KAI2d proteins are involved in chemotropism. Using KAI2 protein sequences found in land plants, we constructed phylogenetic trees. Seven candidate KAI2d homologues encoded in the *P. japonicum* genome^16^ separated into two groups: the first group is similar to KAI2d found in obligate hemiparasites such as *S. hermonthica*, and the second group is similar to those in obligate holoparasites, such as *Orobanche* spp. (Fig. 4a and Supplementary Fig. 4). In combination with transcriptome data in *P. japonicum*^18^, we found that the gene expression levels of *PjKAI2d2*, *PjKAI2d3* and *PjKAI2d3.2,* the *KAI2d* homologues most similar to those in obligate hemiparasites, were high in seedlings. In contrast, little expression was observed in the roots post-infection. This pattern was consistent with the finding that the majority of *KAI2d* genes in *S. hermonthica* were highly expressed in seedlings (Fig. 4a) ^29^. Next, we investigated the effects of *rac*-STR and nutrient conditions on *KAI2* expression in *P. japonicum*. Of the 7 *PjKAI2d* genes as well as *PjKAI2i* and *PjKAI2c*, only *PjKAI2d2* and *PjKAI2d3.2* showed relatively high expression levels in basal conditions and were further increased by *rac*-STR treatment. Basal and induced expression of these genes was attenuated in the nutrient-rich 1/2MS condition (Fig. 4b). Although nutrients also suppressed *PjKAI2d3* expression, the gene was not upregulated by *rac*-STR, unlike *PjKAI2d2* or *PjKAI2d3.2*. The reason why *PjKAI2d3* had a different expression pattern from previous transcriptome data might be because *PjKAI2d3.2* transcripts may have been mistakenly mapped to *PjKAI2d3* due to their high degree of conservation (96% amino acid identity). To test the SL-responding ability of *PjKAI2d2*, *PjKAI2d3* and *PjKAI2d3.2*, we adopted a modified cross-species complementation method, which had been successful in previous studies^27, 43, 44^. We used the *A. thaliana d14 kai2* double mutant in the Col-0 background^45^ and evaluated responses to *rac*-STR by seed germination rates (Fig. 4c). Since Col-0 seeds are known to lose their primary dormancy rapidly after maturation^28^, we stratified seeds at 4 °C overnight after sowing to break dormancy and, therefore, to exclude the effect of dormancy. Germination phenotypes in the control *AtKAI2*-complemented lines were comparable to those in the wild-type Col-0 without significant changes resulting from *rac*-STR treatment, although *D14* was still missing. Concurrently, *rac*-STR promoted germination in *PjKAI2d2-* or *PjKAI2d3.2-*introduced lines with basal germination rates comparable to those of *d14 kai2* (Fig. 4c). These data indicate that at least PjKAI2d and PjKAI2d3.2 are functional as receptors of exogenous SLs.

**Fig. 4:**
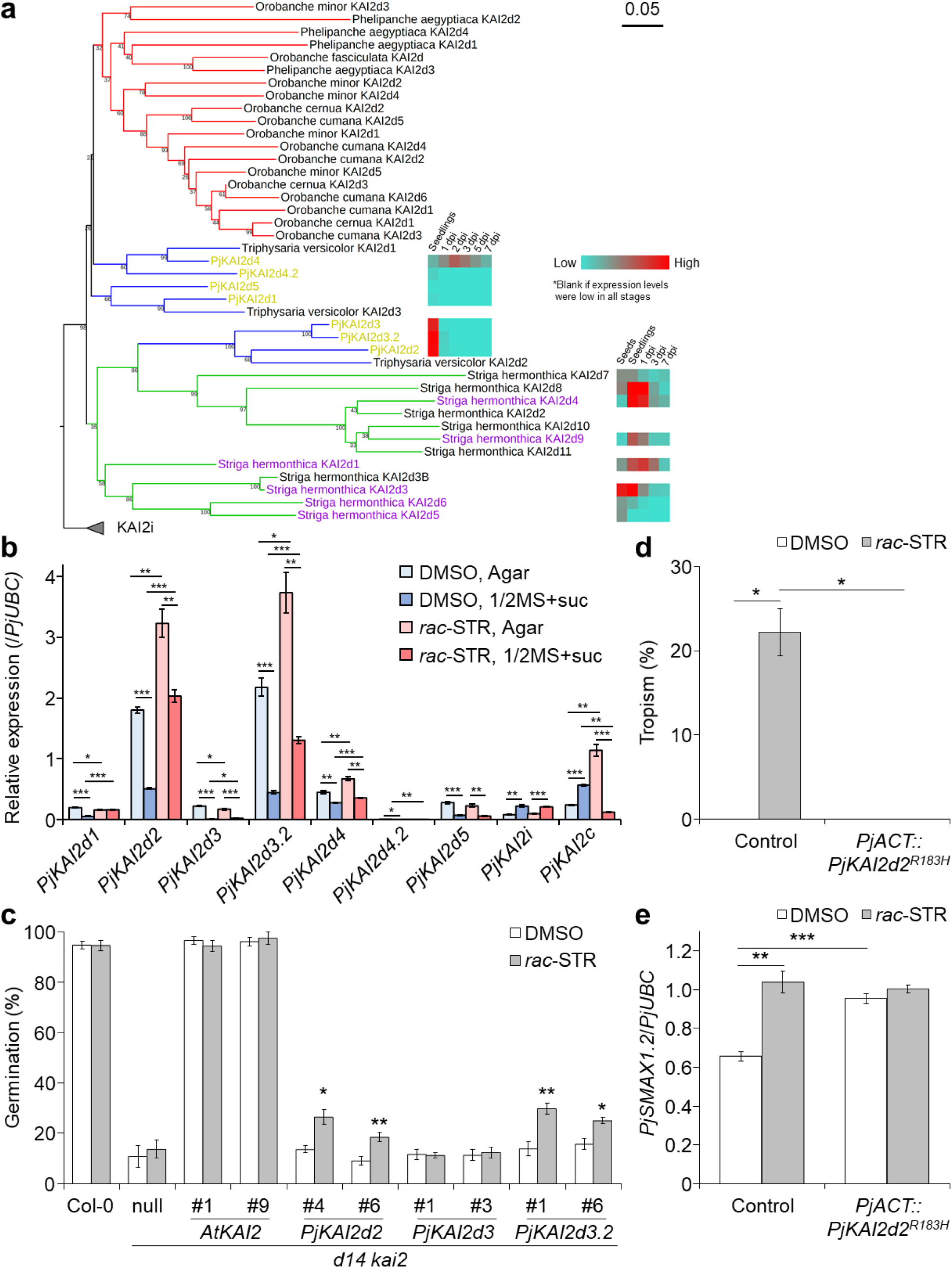
Analyses of KAI2 homologs in *P. japonicum*. **a,** KAI2d phylogeny. The KAI2i clade was included as an outgroup. Clades are coloured blue, green, and red for facultative hemiparasites, obligate hemiparasite, and obligate holoparasites, respectively. Purple and yellow words indicate KAI2d forms in *S. hermonthica* that recognise SLs^30, 72^ and KAI2d forms in *P. japonicum* (PjKAI2d), respectively. Coloured boxes indicate gene expression levels of *PjKAI2d* in roots or *S. hermonthica KAI2d* in seeds, seedlings, or rice plants at 1-, 3-, 7-day post infection (dpi)^18, 29^. The bar indicates substitutions per site. Bootstrap values are indicated at the nodes. **b,** Relative expression levels of *PjKAI2* genes. Representative data are shown (4 technical replicates) using *PjUBC2* as the reference gene. Experiments were performed three times with similar results. **c,** Complementation assays for PjKAI2d. *Arabidopsis kai2 d14* (Col-0 ecotype) was used for the null mutant to express PjKAI2d under control of the *AtKAI2* promoter. Germination rates were calculated 5 days after incubation at 22℃ in the dark. Representative data are shown (4 independent batches, 23-87 seeds per batch). **d-e,** Chemotropic responses to *rac*-STR in *PjKAI2d2* dominant negative plants. (**d**) Percentage of *P. japonicum* transgenic hairy roots that were chemotropic to 1 µM *rac*-STR or 0.1% (v/v) DMSO. Three independent batches (2 to 6 plants) for each compound. Plants that stopped root growth were excluded from the calculations. (**e**) Relative expression level of *PjSMAX1.2*. Representative data are shown (4 technical replicates) using *PjUBC2* as the reference gene. Experiments were performed three times with similar results. **b-e** Mean ± SEM. **b,d,e** *P < 0.05, \*\**P* < 0.01, \*\*\**P* < 0.001 (Welch’s *t* test). **C** \**P* < 0.05, \*\**P* < 0.01, \*\*\**P* < 0.001 (Welch’s *t* test), in comparison to DMSO treatment for each genotype.

To analyse the effects of *KAI2d* genes on chemotropism to SLs, knockout mutants obtained by gene editing would be desirable; however, *KAI2d* genes are multicopy and likely to be functionally redundant (Supplementary Fig. 5). As we cannot yet create transgenerational *P. japonicum* transgenics, it is difficult to edit or knockdown multicopy genes^46^. Hence, we set out to generate dominant-negative plants by overexpressing the substituted KAI2d2, in which the functions of native KAI2d proteins are impeded. KAI2d proteins have a highly conserved arginine that is also conserved in D14 in *A. thaliana* and rice (Supplementary Fig. 5). Substitution of this arginine in D14 of *A. thaliana* and rice with histidine did not interfere with the hydrase activity; however, introducing the substituted D14 into each mutant did not complement the phenotypes due to loss of protein-protein interaction^47^. Therefore, we considered this substitution appropriate for generating KAI2d dominant-negative plants. We transformed *P. japonicum* seedlings with the substituted *PjKAI2d2* (*PjKAI2d2^R183H^*) driven by the constitutively active promoter *PjACT*^17^. Using the resulting transgenic hairy roots, we tested chemotropism to *rac*-STR and quantified the expression levels of *PjSMAX1.2*. Overexpression of *PjKAI2d2^R183H^* hindered chemotropism to *rac*-STR, indicating that *PjKAI2d2^R183H^*-overexpressing hairy roots were dominant negative and that disturbing the interaction of KAI2d with the partner(s) was likely to cause loss of chemotropism to SLs (Fig. 4d). In addition, the expression level of *PjSMAX1.2* was constitutively enhanced in the dominant-negative plants (Fig. 4e), suggesting suppression of SL signalling. Taken together, our results indicate that PjKAI2d proteins play an important role in SL signalling, leading to chemotropism.

## Discussion

Obligate parasitic plants in the Orobanchaceae must attach to host roots within several days after germination to acquire nutrients and water because resources in their seeds are limited, and they cannot obtain sufficient energy for their survival by photosynthesis. Finding a host root to parasitise is also important for facultative parasites, which can survive without a host, as parasitisation significantly enhances its growth^14, 48^. Thus, efficient growth towards host roots is critical for facultative and obligate Orobanchaceae parasites. Our study shows that both facultative *P. japonicum* and obligate *S. hermonthica* use SLs as chemoattractants, leading to host tropism (Fig. 1). As SLs do not induce chemotropism in *L. philippensis*, a non-parasitic member of the Orobanchaceae (Supplementary Fig. 2), this chemotropism phenotype may have been acquired during parasite evolution. Intriguingly, the germ tube of *Orobanche cumana*, a holoparasite, is guided by costunolide, a sunflower-derived sesquiterpene lactone, but not by GR24, a synthetic SL analogue^49^. In comparison, *O. cumana* germination is stimulated by costunolide and GR24^27, 49^. Therefore, it is likely that the nature of chemoattractants is different among phylogenetic clades^2^, in contrast with the commonality of using SLs as germination stimulants in obligate parasites. Host preferences of *Striga* spp.^3^ and *Orobanche* spp.^50^ may be reflected by differences in their chemoattractants. Future identification of specific chemoattractants produced by each host may uncover the chemical basis for host preference in Orobanchaceae parasites.

SLs are known to stimulate hyphal branching of arbuscular mycorrhizal fungi (AMF)^51^ that establish symbiotic relationships with plants to exchange soil-derived nutrients such as phosphate and nitrogen with plant-derived carbon sources^52, 53^. Interestingly, D14L, a KAI2 homologue, is required for AMF symbiosis in rice^54^. D14L derepresses downstream signalling by removing the suppressor SMAX1, resulting in elevated AMF colonization^39^. In our study of *P. japonicum*, we found that disturbance of KAI2d function(s) by overexpressing a substituted KAI2d2 resulted in elevated *PjSMAX1.2* expression and loss of chemotropism to SLs (Fig. 4d,e). As KAI2d-mediated SL perception for chemotropism is important in host infection to obtain nutrients, it is tempting to think that Orobanchaceae parasites may have converted the KAI2-based symbiotic AMF communication tool via MAX2 into a parasitic host-detection system. In this context, the observation that chemotropism to SLs occurs only in nitrogen-deficient conditions is intriguing (Fig. 2). Plants often produce and exude SLs when nitrogen is deficient for attracting AMF^55^, which can, in turn, promote plant nitrogen acquisition. Indeed, in the case of rice, ∼40% of the plant’s nitrogen requirement can be acquired by AMF^56^. It is possible that, in Orobanchaceae parasites, the primary nitrogen source may have shifted from AMF to host plants, potentially by evolving KAI2-mediated signalling via MAX2^57^. Consistently, *P. japonicum* effectively transfers nitrogen from hosts, especially in nutrient-deficient conditions^58^. AMF colonisation in *P. japonicum* has not been reported, but some *Pedicularis* species, facultative hemiparasitic plants in the Orobanchaceae, can accommodate AMF^59^. How such parasites can coordinate SLs and KAI2/KAI2d proteins to accommodate both AMF and hosts is an interesting question to be answered. Future genomic surveys of the AMF-accommodating abilities in Orobanchaceae plants may provide molecular clues of how the KAI2/KAI2d-based symbiotic-parasite relationship has evolved.

Complementation of the *d14 kai2* mutant with *PjKAI2d2* or *PjKAI2d3.2* partially rescued seed germination in an exogenous SL-dependent manner (Fig. 4c). This result indicates that SL-bound PjKAI2d2 and PjKAI2d3.2 may not efficiently bind to *A. thaliana* MAX2 and/or SMAX1. In addition, *PjKAI2d2* and *PjKAI2d3.2* failed to complement the germination-attenuated phenotype without SL application, indicating that these proteins are unlikely to bind to unidentified endogenous KAI2 ligands (KLs) to activate downstream signalling. Similarly, *S. hermonthica KAI2c*, *KAI2i*, *KAI2d1* and *KAI2d2* do not complement the *A. thaliana kai2* mutant without SLs^27^, suggesting that *S. hermonthica* may have lost KL-sensing activity for germination. Further characterisation of various KAI2 proteins and associated proteins in the Orobanchaceae family and other land plants will reveal signalling specificities differentiating chemotropism and germination functions. This research direction may eventually provide clues of how neo-functionalisation of KAI2/KAI2d occurs in plants.

Our study shows that positioning SLs beside *P. japonicum* roots induces asymmetrical accumulation of the auxin efflux transporter PIN2, resulting in the roots bending towards the chemicals (Fig. 3). This observation provides a functional link between SL and auxin signalling in the host tropism of *P. japonicum*. To date, the effects of SLs on auxin-mediated processes have been studied primarily in *A. thaliana*^60, 61^. SLs interfere with auxin-mediated PIN polarisation in *A. thaliana* in a MAX2-dependent manner by derepressing PIN endocytosis, resulting in the interference of auxin canalisation^62^. In roots, asymmetrical PIN2 accumulation in epidermal cells is a key factor in root bending, as seen in gravitropism^62^. In *P. japonicum*, exogenous SLs are perceived by the epidermal cells in the elongation zone, as indicated by YLG-based fluorescence, where PjPIN2 accumulates (Fig. 1i and 3n). This result is consistent with the finding that KAI2 is required for the local accumulation of PIN2 in *A. thaliana* roots^63^. As *A. thaliana* does not exhibit chemotropism to SLs (Supplementary Fig. 2c,d) and frequently SLs are evenly distributed throughout *A. thaliana* media, it is difficult to compare the effects of SLs on PIN2 accumulation in *A. thaliana* and *P. japonicum*. An alteration in SL-PIN2 relationships may have occurred in Orobanchaceae parasites that enabled chemotropism. Importantly, SLs can be perceived in nutrient-rich conditions (Fig. 2e), but the asymmetrical PIN2 accumulation occurs only in nitrogen-, especially ammonium ion-, deficient conditions (Fig. 3n and 3p). The SL signalling pathway is also affected by ammonium ions, as indicated by upregulation of *SMAX1.1* and *SMAX1.2* (Fig. 2i-k). This finding suggests that ammonium ions regulate root tropism by controlling PIN2 accumulation potentially via SMAX regulation. As rice roots exhibit chemotropism to ammonium ions^64^, it would be interesting to test whether that tropism to ammonium ions also involves an asymmetric accumulation of PIN2, which may be regulated by SL signalling in rice.

In summary, our study unveils important molecular clues about host tropism in Orobanchaceae parasitic plants, providing a novel function of SLs as chemoattractants. We expect this study will encourage additional investigations to elucidate tropism processes, one of the important steps for infection in parasitic plants. Such future studies will help design solutions for protecting agricultural fields from nuisance weeds.

## Methods

### Plant materials and growth conditions

Unless otherwise noted, *P. japonicum* (Thunb.) Kanitz seeds and *A. thaliana* seeds (Col-0, *d14 kai2*^45^ and complemented lines) were germinated on 1/2MS medium (0.8% (w/v) INA agar, pH 5.8) containing 1% (w/v) sucrose. Before sowing, seeds were sterilised with a diluted commercial bleach solution (Kao, Tokyo, Japan, 10% (v/v) for *P. japonicum* and 5% (v/v) for *A. thaliana*) for 5 minutes and rinsed at least 5 times with sterilised water. Plate-sown seeds were stratified at 4 °C in the dark for 1 to 3 night(s), then grown horizontally in long-day conditions (16-h light (∼40 and ∼30 μmol m^-2^s^-1^ for *P. japonicum* and *A. thaliana*, respectively), 8-h dark at 25 °C and 22 °C for *P. japonicum* and *A. thaliana*, respectively). *P. japonicum* and *A. thaliana* seeds were grown at 70% and 50% humidity, respectively. *S. hermonthica* seeds were carefully handled with glass pipettes as the seeds stick to plastic pipette tips. Seeds were soaked in a diluted 20% (v/v) commercial bleach solution during a brief vortex, then sterilised with a fresh bleach solution for 5 minutes, followed by at least 5 rinses with sterilised water. Seeds were then soaked in 5-mL of sterilised water in 6-well plates and incubated in the dark at 25 °C for 1 to 2 weeks. To induce germination, water was replaced with a 10 nM (+)-strigol (STR)^65^ solution, and the plates were incubated for 4 hours in the dark at 25 °C. The (+)-STR solution was then replaced with water, and the plates were incubated at least 24 hours before further analysis to exclude the residual effects of (+)-STR. *L. philippensis* seeds were sterilised with a diluted commercial bleach solution (Kao, Tokyo, Japan, 5% (v/v)) for 5 minutes and rinsed at least 5 times with sterilised water before soaking in 5-mL of sterilised water in 6-well plates. Water-soaked seeds were grown in long-day conditions (∼30 μmol m^-2^s^-1^) at 22 °C for six days. For further analyses, seedlings were carefully handled with glass pipettes and a pair of tweezers. *O. sativa* (japonica, c.v. Shiokari) seeds of WT and *d10*^35^ were sterilised with 70% (v/v) ethanol for 3 minutes, followed by a diluted commercial bleach solution (Kao, Tokyo, Japan, 50% (v/v)) for 30 minutes and rinsed with sterilized water at least 5 times. Surface-sterilized seeds were then grown vertically on 0.6% (w/v) water agar in long-day conditions (18 h /6 h light/dark, ∼120 μmol m^-2^s^-1^) at 25 °C.

### Chemicals

SLs and analogues used in this study are listed in Supplementary Table 1. Chemicals were stored at - 20 °C as 100 mM (YLG) or 10 mM (the other chemicals) stocks in DMSO.

### Chemotropism assays

For assays using *P. japonicum* seedlings, 3-day-old seedlings were transferred carefully to solid media containing nutrients (0.8% (w/v) INA agar, pH 5.8) or solid media without nutrients (0.7% (w/v) INA agar, pH 5.8), incubated vertically for 1 day, and transferred to the media used for the assays. Filter paper disks (4-mm diameter) were soaked with the chemical-containing solution before being placed 5-mm from the seedlings. *P. japonicum* transgenic hairy roots were generated from seedlings 3- to 4- weeks post-transformation and were transferred carefully to solid medium without nutrients (0.7% (w/v) INA agar, pH 5.8), incubated horizontally for 2 days, and transferred to new solid medium without nutrients. Filter paper disks (4-mm diameter) were soaked with a chemical-containing solution before being placed 5-mm from the seedlings. For assays using *S. hermonthica*, germinated seedlings were transferred carefully to solid medium without nutrients (0.7% INA agar (w/v), pH 5.8). Filter paper disks (4-mm diameter) were soaked with a chemical-containing solution, before being placed 3- mm from the seedlings. For assays using *L. philippensis*, germinated seedlings were transferred carefully to solid medium without nutrients (0.7% INA agar (w/v), pH 5.8). Filter paper disks (4-mm diameter) were soaked with a chemical-containing solution, before being placed 3-mm from the seedlings. For assays using *A. thaliana*, 2-day-old seedlings were transferred carefully to solid medium without nutrients (0.7% (w/v) INA agar, pH 5.8), incubated vertically for 1 day, and transferred to new solid medium. Filter paper disks (4-mm diameter) were soaked with a chemical-containing solution, before being placed 5-mm from the seedlings. We defined seedlings that exhibited chemotropism as showing root bending of 30° or more towards a filter paper disk containing a tested chemical.

### Microscopy

For fluorescence microscopy, photos were taken using a fluorescence stereo microscope (M165 FC, Leica) before and after chemical-treated seedlings were incubated in the dark at 70% humidity for 1 day, 25 °C for *P. japonicum* and *S. hermonthica*, and 22 °C for *A. thaliana* and *L. philippensis*, respectively, except for time-lapse microscopy. For time-lapse microscopy, 100 photos were taken throughout 18 hours using a fluorescence stereo microscope (M205 FA, Leica) in the dark at 25 °C. For confocal microscopy, photos were taken using an inverted confocal microscope (TCS SP5 II, Leica). Fluorescein from hydrolysed YLG was excited with a 488-nm laser, and the detected emission spectra were observed at 500–531nm. Excitation and detection of Venus and mCherry fluorescence were performed as described previously^12^. The bending degree and image analyses were measured using ImageJ^66^.

### Host chemotropism assay for *P. japonicum*

*P. japonicum* seeds were sterilised, sown, stratified, and grown on solid growth medium containing 1/2 MS, 0.6% (w/v) agar, and 1% sucrose (pH 5.8) for 3 days in the dark at 25 °C. The seedlings were then transferred to 0.6% (w/v) agar medium without nutrients and grown vertically for 1 day in long-day conditions at 25 °C. Rice seeds were sterilised, sown, and grown for 4 days as described in the *Plant materials and growth conditions* section. Host chemotropism assays were then performed on 0.6% agar medium without nutrients in long-day conditions at 25 °C as follows: a pair of WT rice and *d14* rice seedlings were transferred to agar medium in a square petri dish (140 mm x 100 mm x 15 mm). Single roots of similar length from each genotype were chosen and aligned vertically in parallel at a distance of approximately 2 mm. A *P. japonicum* seedling was then placed between two rice roots and a single *P. japonicum* root was carefully placed parallel to the middle of the rice roots. Relatively young root regions (approximately 500 mm from the tip) of the host were chosen for the initial infection position (0 days) because these areas produce few to no lateral roots during the observation period that would otherwise physically disturb *P. japonicum* roots from bending. Three independent experiments were conducted with each replicate containing 7 pairs. For about half of the pairs, WT seedlings were placed at the left and *d10* seedlings at the right, and for the rest vice versa. To keep the roots on the medium, a cover glass was placed on the top of the roots. The plates were sealed with surgical tape and positioned vertically in the growth chamber. Images were captured 0, 1, and 2 day(s) after infection using a wide zoom stereo microscope (Olympus SZX16).

ImageJ (version 1.52q), Microsoft Excel and Adobe Illustrator software were employed to quantify the images. In all cases, WT roots were positioned on the left and *d10* was on the right. First, images derived from 1- or 2-day(s) post-infection were superimposed onto their corresponding 0-day images using Adobe Illustrator. Next, using ImageJ, the position of the *P. japonicum* root tip was defined as y = 0 and the position of the left (WT) host at horizontal axis with the *P. japonicum* root tip was defined as x = 0 in 0-day samples. Thus, each rice root and the *P. japonicum* root tip were assigned a unique x and y value: for instance, WT rice (x = 0 µm, y = 0 µm), *P. japonicum* (x = 904.8 µm, y = 0 µm) and *d10* rice (x = 1979.6 µm, y = 0 µm) at 0 day. The xy positions of the *P. japonicum* root tip at 1- and 2-day(s) post infection relative to 0 day were then measured. Raw xy data generated from all samples as described above were subjected to centring and scaling processes before generating the graphs shown in the figures (For details refer to Source Data).

### Extraction of total RNA and RT-qPCR

To extract total RNA from *P. japonicum* seedlings, 3-day-old seedlings were transferred to media containing varying nutrients and were grown vertically for 1 day. The seedlings were transferred again to the same medium used for the assays. At 3 hours after incubation, root tips were excised and immediately frozen in liquid nitrogen. To extract total RNA from *P. japonicum* transgenic hairy roots, root tips of the generated hairy roots were excised and immediately frozen in liquid nitrogen after the chemotropism assays (see *Chemotropism assay* section). Total RNA extraction, cDNA synthesis and RT-qPCR were performed as previously described^12^.

### Cloning

The constructs *pDR5::3xmCherry-NLS* and *pPjPIN2::PjPIN2-Venus*^19^ were previously described. pKAI2pro-GW^67^ was used as the Gateway destination vector to express each protein under the control of the *AtKAI2* promoter. The AtKAI2-expressing vector pKAI2-AtKAI2 was previously described^67^. PjKAI2d-expressing vectors were constructed as follows: the genomic regions containing each coding sequence (CDS) of *PjKAI2d* was PCR amplified. With the resulting PCR products, each PjKAI2d CDS was PCR amplified with primers containing attB sites, subcloned into pDONR/Zeo (Thermo Fisher Scientific) by Gateway BP cloning (Thermo Fisher Scientific) to generate entry vectors, then cloned into pKAI2pro-GW by Gateway LR cloning (Thermo Fisher Scientific), yielding pKAI2-PjKAI2d2, pKAI2-PjKAI2d3 and pKAI2-PjKAI2d3.2. For constructing the expression vectors to transform *P. japonicum*, we used GoldenGate technology for assembling modules^68^. The GoldenGate modules containing the *35S* promoter (*p35S*, pICH51266) or *35S* terminator (*35St*, pICH41414) were pre-existing^68^. Modules containing the actin promoter (*pPjACT*), 3xVenus-NLS CDS or HSP18.2 terminator with the 3ʹ-untranslated region (*HSPt*) were described previously^13, 17^. A single mutation was introduced into the *PjKAI2d2* CDS using a KOD plus Mutagenesis kit (TOYOBO) with the entry vector described above as a template. The mutated *PjKAI2d2* CDS was PCR amplified and cloned into pICH41308 to generate a level 0 CDS1 module. The modules were assembled and cloned into level 1 vectors to generate *pPjACT::PjKAI2d2^R183H^:HSPt* and *p35S::3xvenus-NLS:35St* transcription units. The resulting transcription units were further combined and cloned into the level 2 binary vector pAGM4723. End-linkers and dummies were used as needed.

### Transformation of *P. japonicum*

*Agrobacterium rhizogenes* AR1193 strain was used to transform *P. japonicum* seedlings. *A. rhizogenes*-mediated transgenic hairy roots were generated and identified according to previously described methods^12, 13^.

### Phylogenetic analyses

We used the CLC Main Workbench (ver. 8.0, Qiagen) for phylogenetic analyses. The CDSs of KAI2 used for the phylogenetic analyses in Conn *et al.*^27^ were automatically translated and used in our study. Two candidate genes, KAI2d3.2 and KAI2d4.2 in *P. japonicum*, which were newly identified from the *P. japonicum* genome by BLASTp analysis using PjKAI2d1-PjKAI2d5 as queries (with e-values under 1e^-100^), were added to the KAI2 sequence group. The KAI2 sequences processed by trimAL v1.2^69^ using the automated1 settings were aligned using default settings. After alignment, phylogenetic trees were drawn using the maximum-likelihood method with 1,000 bootstrap repetitions. We generated the figures using iTOL v6 (https://itol.embl.de/ ^70^).

### Transformation of *A. thaliana* and cross-species complementation assays

pKAI2-AtKAI2, pKAI2-PjKAI2d2, pKAI2-PjKAI2d3 or pKAI2-PjKAI2d3.2 were electroporated into *Agrobacterium tumefaciens* GV3101 competent cells. The resulting bacterial cells were used to inoculate flowers in a transformation method modified from Martinez-Trujillo *et al.*^71^. For germination assays, *A. thaliana* seeds that were at least 1-month old were used. Seeds were sterilised with chlorine gas for 2–3 hours in a safety cabinet and dried for at least 2 hours in a clean bench, followed by sowing on solid media without nutrients (0.7% (w/v) INA agar, pH 5.8) containing 1 µM *rac*-STR or 0.1% (v/v) DMSO. Plate-sown seeds were stratified at 4 °C in the dark for 1 night, then incubated horizontally in the dark. Germination rates, defined by radicle emergence, were scored periodically.

### Statistical analyses

Welch’s *t* test and two-way ANOVA, Tukey’s multiple comparison test were performed in Microsoft Excel 2016 and GraphPad Prism version 8.1, respectively. Details of statistical analyses including statistical methods, numbers of individual batches, numbers of technical replicates, numbers of biological replicates, and statistical significances are described in each figure legend.

## Data and code availability

Transcriptome data for *S. hermonthica*^29^ and *P. japonicum*^18^ are available from the DNA Data Bank of Japan (http://www.ddbj.nig.ac.jp/) under accession numbers DRA008615 and DRA003608 for *S. hermonthica* and DRA010010 for *P. japonicum*, respectively. Sequence data from this study have been deposited at the GenBank/EMBL libraries and are publicly available as of the date of publication. Accession numbers are provided in Supplementary Table 2. Gene IDs of the *P. japonicum* genes investigated in this study are provided in Supplementary Table 2. Source data are provided with this manuscript.

## Acknowledgements

We thank Prof. Yuichiro Tsuchiya (Institute of Transformative Bio-Molecules, Nagoya University), Prof. Lam-Son Phan Tran (Institute of Genomics for Crop Abiotic Stress Tolerance, Texas Tech University) and the late Prof. Kenji Mori for sharing *L. philippensis* seeds, *A. thaliana d14 kai2* mutant seeds and (+)-strigol, respectively. This work was supported by Ministry of Education, Culture, Sports, Science and Technology KAKENHI grants (19K16169 to S.C., 20H05909 and 21H02506 to K.S. and S.Y., 17H06172 to K.S); the Japan Society for the Promotion of Science (JSPS) KAKENHI Grants-in-Aid for JSPS Fellows (JP21J00718 to S.O.); JST PRESTO (JPMJPR194D to S.Y.).

## Author Contributions

S.O., D.C.N., S.Y., and K.S. conceived and designed the study; K.S. supervised the experiments; S.O., the original manuscript; all authors critically revised the manuscript and approved the final version.

## Ethics declaration

The authors declare no competing interests.

## Additional information Supplementary information

Supplementary Figure 1: **Chemical structure of SLs and synthetic analogues used in this study.**

Supplementary Figure 2: **Chemotropic response to *rac*-STR and YLG in non-parasitic plants.**

Supplementary Figure 3: **Chemotropic phenotype to *rac*-STR on nutrient-containing media.**

Supplementary Figure 4: **Classification of KAI2 proteins in dicots.**

Supplementary Figure 5: **Alignment of KAI2d, KAI2i and KAI2c proteins in *P. japonicum*, KAI2d in *S. hermonthica*, KAI2 and D14 in *A. thaliana*, and D14 and D14L in rice.**

Supplementary Table 1: **Chemicals used in this study.**

Supplementary Table 2: **Gene IDs and accession numbers of the *P. japonicum* genes investigated in this study.**

Supplementary Table 3: **Primers used in this study.**

Supplementary Video 1 **Time-lapse imaging of chemotropism to YLG in *P. japonicum*.**

Supplementary Video 2 **Time-lapse imaging of tropism to rice in *P. japonicum*.**

Supplementary Video 1 **Time-lapse imaging of chemotropism to YLG in *P. japonicum*.** Filter paper disks were soaked in 1 µM YLG (left) or 0.1% (v/v) DMSO (right) and placed 5-mm from the root of a 3-day old *P. japonicum* seedling growing on 0.7% INA agar medium (w/v). Images were taken for 18 hours at intervals of 648 seconds.

Supplementary Video 2 **Time-lapse imaging of tropism to rice in *P. japonicum*.** Root growth of *P. japonicum* was captured for 48 hours at intervals of 20 minutes after being placed between the wild-type rice (top) and *d10* (bottom) seedlings growing on 1% water agar medium (w/v). Video was generated with 12 frames per second.

## Notes

### Competing Interest Statement

The authors have declared no competing interest.

