## Supplementary Figures for "Strigolactones are chemoattractants for host tropism in Orobanchaceae parasitic plants"

**Strigolactones**

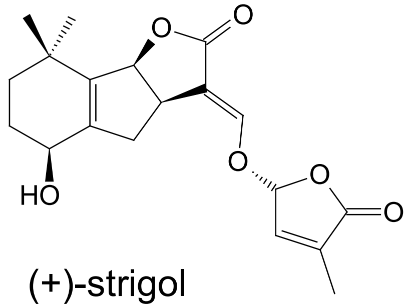

(+)-strigol

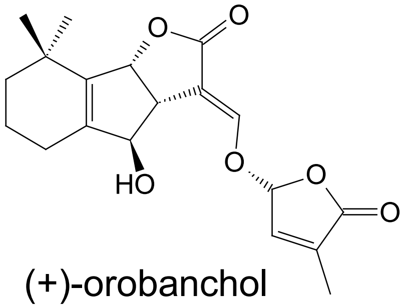

(+)-orobanchol

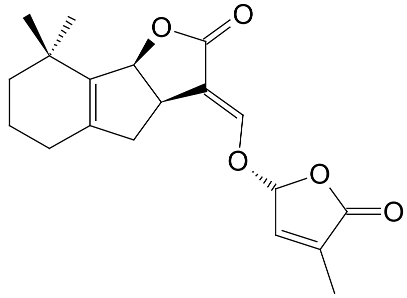

(+)-5-deoxystrigol

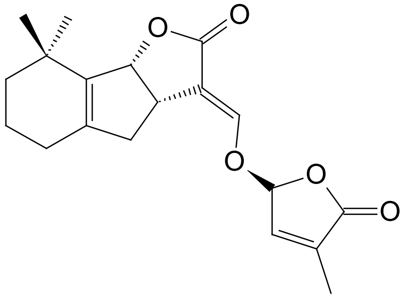

(-)-5-deoxystrigol

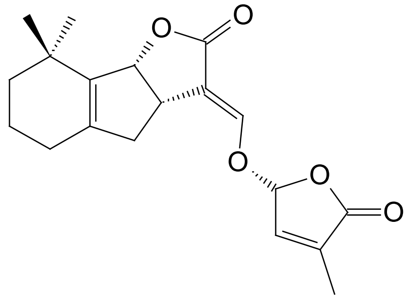

(+)-2’-*epi*-5-deoxystrigol

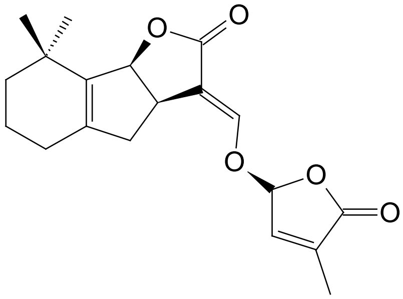

(-)-2’-*epi*-5-deoxystrigol

**Synthetic analogues**

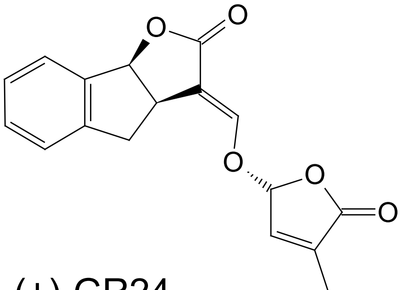

(+)-GR24

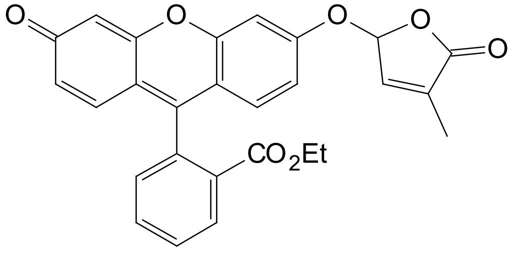

Yoshimulactone green

Supplementary Figure 1: **Chemical structure of SLs and synthetic analogues used in this study.** (+) forms are indicated for strigol, orobanchol and GR24.

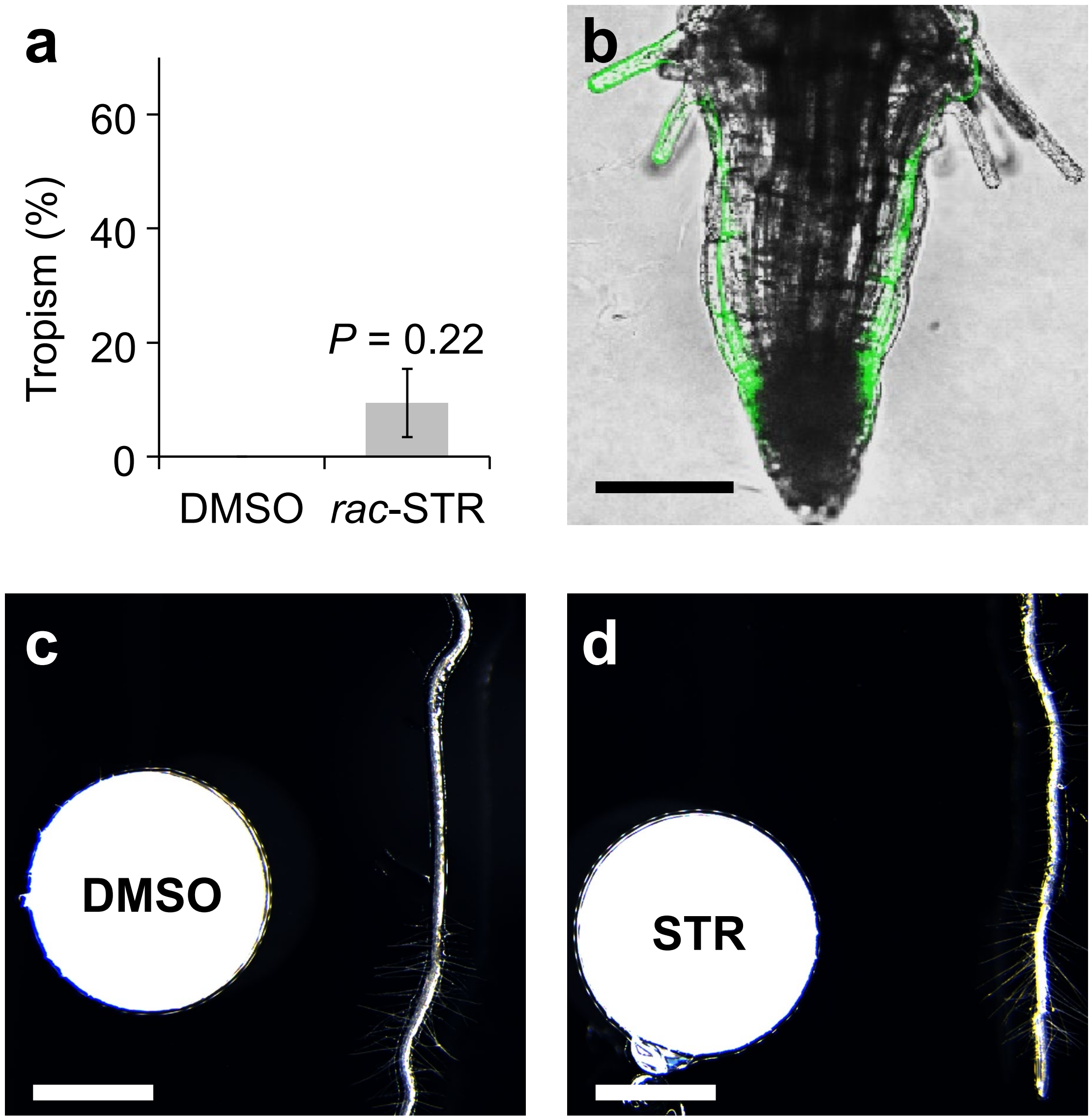


Supplementary Figure 2: **Chemotropic response to *rac*-STR and YLG in non-parasitic plants.**
**a,** Ratio of *L. philippensis* plants that showed chemotropism to 1 μM *rac*-STR or 0.1% (v/v) DMSO. Three or four independent batches for each compound, 4-8 plants tested per batch. Plants that stopped root growth were excluded from ratio calculation. **b,** A representative image of *L. philippensis* plant that showed YLG-derived fluorescence when treated with a 100 μM YLG solution. Filter paper disks were placed 3-mm to the left of the roots. Confocal photos were taken 24 hours after treatment. Bar = 100 μm. **c and d,** Representative images of *A. thaliana* plants tested for chemotropism. Photos were taken 1 day after treatment. c, 0.1% (v/v) DMSO; d, 1 μM *rac*-STR. At least 18 plants were tested for each chemical with similar phenotypes. Bars = 1 mm.

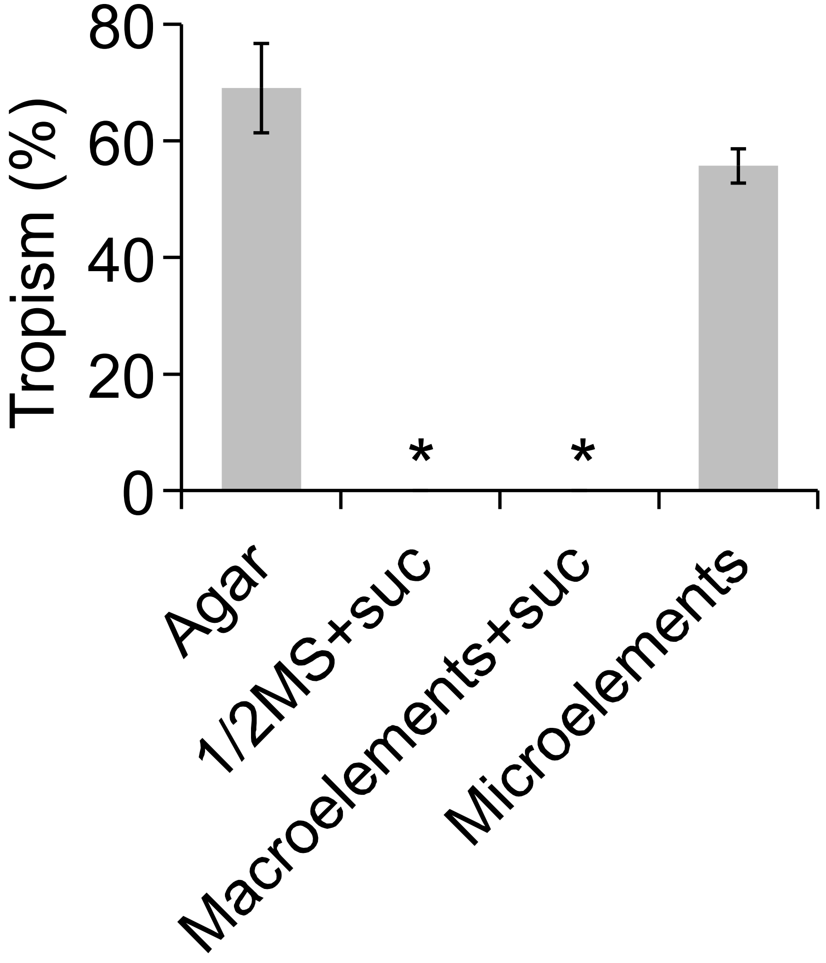


Supplementary Figure 3: **Chemotropic phenotype to *rac*-STR on nutrient-containing media.**
Ratio of *P. japonicum* plants that showed chemotropism to 1µM *rac*-STR in each medium. Three independent batches for each compound, 6 to 8 plants tested per batch. Plants that stopped root growth were excluded from ratio calculation. Asterisks indicate statistical significance in comparison with no nutrient condition (Welch's *t* test, $^{*}P < 0.05$)

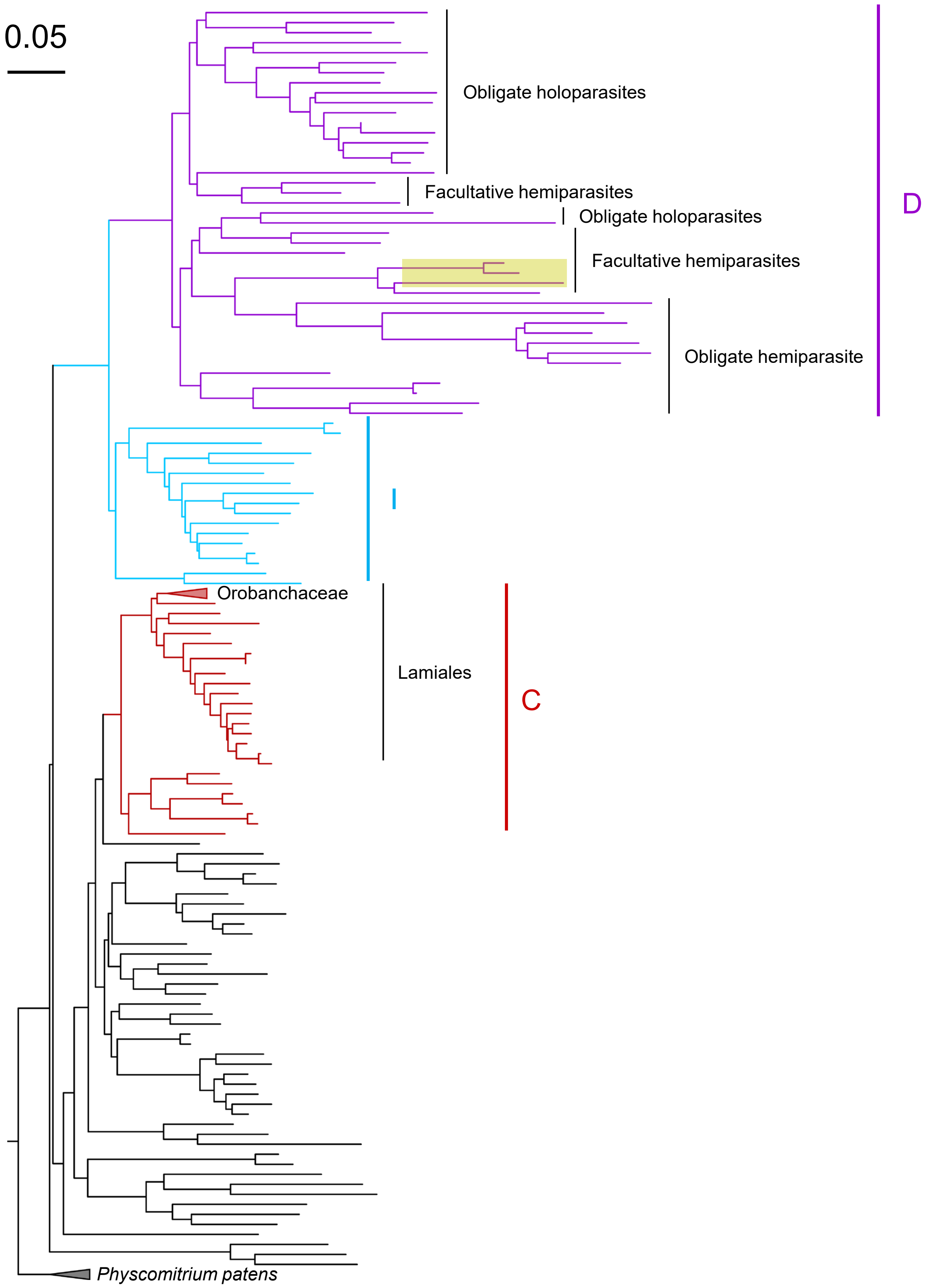


Supplementary Figure 4: **Classification of KAI2 proteins in dicots.**
*Physcomitrium patens* KAI2 was included as an outgroup. Clades in the Lamiids were colored red, blue and violet for conserved (C, KAI2c), intermediate (I, KAI2i), and divergent (D, KAI2d), respectively. Orobanchaceae clade in the KAI2c was collapsed. PjKAI2d2, PjKAI2d3, PjKAI2d3.2 were highlighted in yellow. A Bar indicates substitutions per site.

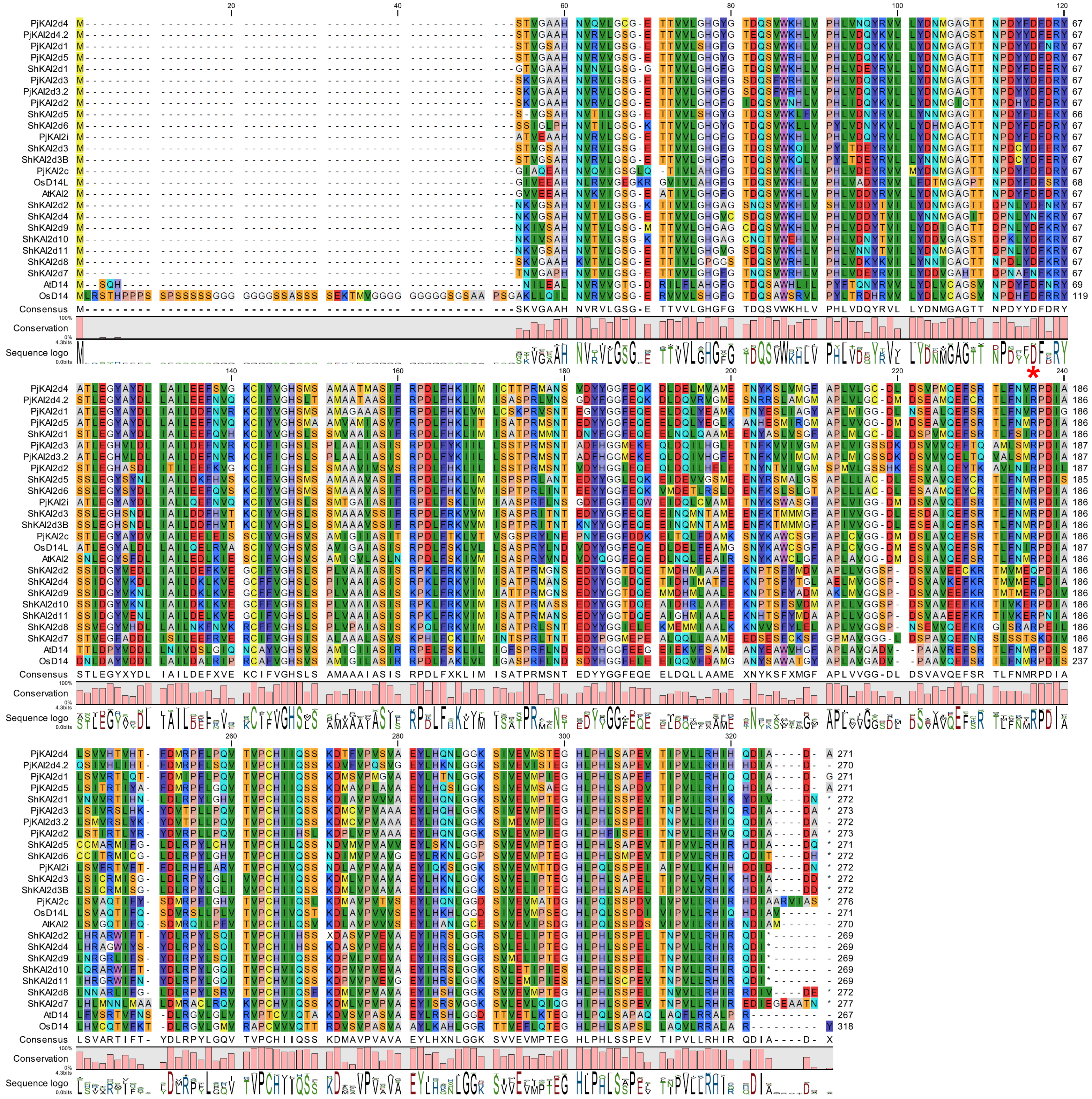

Supplementary Figure 5: **Alignment of KAI2d, KAI2i and KAI2c proteins in *P. japonicum*, KAI2d in *S. hermonthica*, KAI2 and D14 in *A. thaliana*, and D14 and D14L in rice.** A red asterisk indicates the substituted residue of PjKAI2d for dominant negative.
